# Kilohertz-rate two-photon voltage imaging of population dynamics *in vivo*

**DOI:** 10.64898/2025.12.29.696942

**Authors:** Min Zhang, Shoupei Liu, Yaoguang Zhao, Yanfeng Zhu, Xinyang Gu, Cihang Kong, Jiahao Hu, Huiyun Yu, Jingchuan Wu, Fang Xu, Liang Chen, Ying Mao, Bo Li

**Affiliations:** Jinshan Hospital, MOE Frontiers Center for Brain Science, Institutes for Translational Brain Research, Fudan University, Shanghai 200032, China; School of Psychology, Institute of Chinese Cultural Psychology, Shanghai Jiao Tong University, Shanghai 200030, China; Department of Neurosurgery, Huashan Hospital, Shanghai Medical College, Fudan University, Shanghai 200040, China; National Center for Neurological Disorders, Shanghai, China; Department of Pharmacology, School of Basic Medical Sciences, Fudan University, Shanghai 200032, China; Interdisciplinary Center for Brain Information, Brain Cognition and Brain Disease Institute, Shenzhen Institutes of Advanced Technology, Chinese Academy of Sciences, Shenzhen, 518055, China; Jinshan Hospital, Huashan Hospital, Institutes for Translational Brain Research, Fudan University, Shanghai 200032, China

## Abstract

Understanding how neural circuits compute requires capturing voltage dynamics across large neuronal populations with millisecond resolution *in vivo*. However, two-photon voltage imaging remains fundamentally limited by a trade-off among imaging speed, field of view, and excitation efficiency. We introduce HS2PM, a hybrid scanning two-photon microscope that overcomes this bottleneck, enabling kilohertz-rate imaging over a 650 × 524 μm^2^ field of view at single-cell resolution while preserving the photon efficiency of single-point excitation. HS2PM stably records deep-layer membrane potentials from over 160 neurons simultaneously, with minimal photobleaching and phototoxicity. It resolves both spikes and subthreshold voltage dynamics *in vivo*, revealing how these jointly shape sensory adaptation and population coding. Beyond voltage imaging, HS2PM supports high-speed fluorescence lifetime and vascular flow imaging, establishing a multimodal platform for dissecting fast circuit dynamics with precision previously inaccessible to optical methods.

## Introduction

Neural circuits compute at millisecond timescales, coordinating rapid voltage changes across distributed neuronal populations. Capturing these fast dynamics *in vivo* is essential for understanding how brain activity underlies behavior^1,2^. While calcium imaging has revolutionized population-scale neural recording^3,4^, its inherently slow kinetics obscure the fast electrical signals that drive synaptic integration and action potentials^2,5,6^.

Two-photon voltage imaging (TPVI) with genetically encoded voltage indicators (GEVIs) offers a path toward high-speed, deep-tissue recordings of membrane potential^7–13^. However, efforts to scale TPVI to large populations remain constrained by a three-way trade-off between imaging speed, field of view (FOV), and excitation efficiency. Parallelized approaches such as multi-point^7,14^, line^15^, or light-sheet^16,17^ scanning increase acquisition rates by distributing excitation across space, but they sacrifice photon efficiency, increase laser power demand^18^, and introduce cross-talk in scattering tissue^7^. In contrast, single-point scanning maximizes signal-to-noise ratio (SNR) and the number of sampled neurons^18–20^, but it is limited by the speed of mechanical deflection systems^21–23^.

To overcome these limitations, we developed HS2PM, a hybrid scanning two-photon microscope that integrates high-speed scanning with photon-efficient single-point excitation. This system achieves kilohertz-rate imaging across a 650 × 524 μm^2^ FOV at single-cell resolution. HS2PM enables stable, high-SNR recordings from over 160 neurons in awake mice, reaching depths of 700 µm with minimal phototoxicity. It resolves both spikes and subthreshold voltage dynamics *in vivo*, revealing how they jointly contribute to sensory adaptation and population coding. In addition to voltage imaging, HS2PM supports high-speed fluorescence lifetime and vascular flow imaging, offering a versatile platform for multimodal interrogation of circuit dynamics at millisecond resolution.

## Results

### Overcoming the speed-efficiency bottleneck

TPVI is fundamentally limited by a trade-off between excitation efficiency and imaging speed. Single-point scanning provides high efficiency but limited speed, whereas parallel strategies improve speed at the expense of per-focus efficiency and fidelity in scattering tissue. This efficiency loss is particularly detrimental for voltage imaging, where the photon budget is already tight due to membrane-localized GEVIs and millisecond-scale dynamics. A theoretical analysis (Supplementary Note 1) showed that, even under ideal single-point conditions, voltage imaging would require more than 50-fold greater laser power than calcium imaging to achieve comparable discriminability index (*d’*). When parallel strategies further divide excitation across multiple foci, the effective power requirement multiplies^18^, rapidly driving the system beyond safe limits (≤250 mW at 920 nm^24^). Together, these considerations establish that voltage imaging fundamentally requires single-point excitation to maximize efficiency, while speed gains must instead be realized through innovations in scanning architecture.

HS2PM overcomes the speed bottleneck of single-point scanning, achieving kilohertz-rate imaging across a large population-scale FOV. This is enabled by a hybrid scanning architecture that combines fast angular encoding with high line-rate deflection (Fig. 1a; Extended Data Fig. 1). First, an electro-optic deflector (EOD) angularly encoded the 80 MHz pulse train, cycling every four pulses to generate four spatiotemporally indexed points (Fig. 1b). These were further expanded by a spatiotemporal division multiplexing (STDM) module, which split each pulse into four time-staggered replicas, yielding 16 spatiotemporally indexed points at an effective sampling clock of 320 MHz (Fig. 1c). A 36-facet polygon scanner rotating at 54,945 rpm then provided a base line rate of ∼33 kHz. Scanning the 16 indexed points generated 16 lines, with an effective line rate of 527 kHz (Fig. 1d). Finally, a bidirectionally driven galvo performed *y*-axis scanning to generate 544 lines per frame (Fig. 1e). Together, this architecture enabled 916 Hz across 571 × 544 pixels (650 × 524 μm^2^ FOV; Fig. 1f; see also Extended Data Fig. 2 for pixel order map). Among previously reported fast point-scanning systems capable of voltage imaging^11,20,25–27^, HS2PM achieves the largest FOV and the highest sampling speed at kilohertz frame rates (Fig. 1g; Supplementary Table 1).

**Fig. 1.**
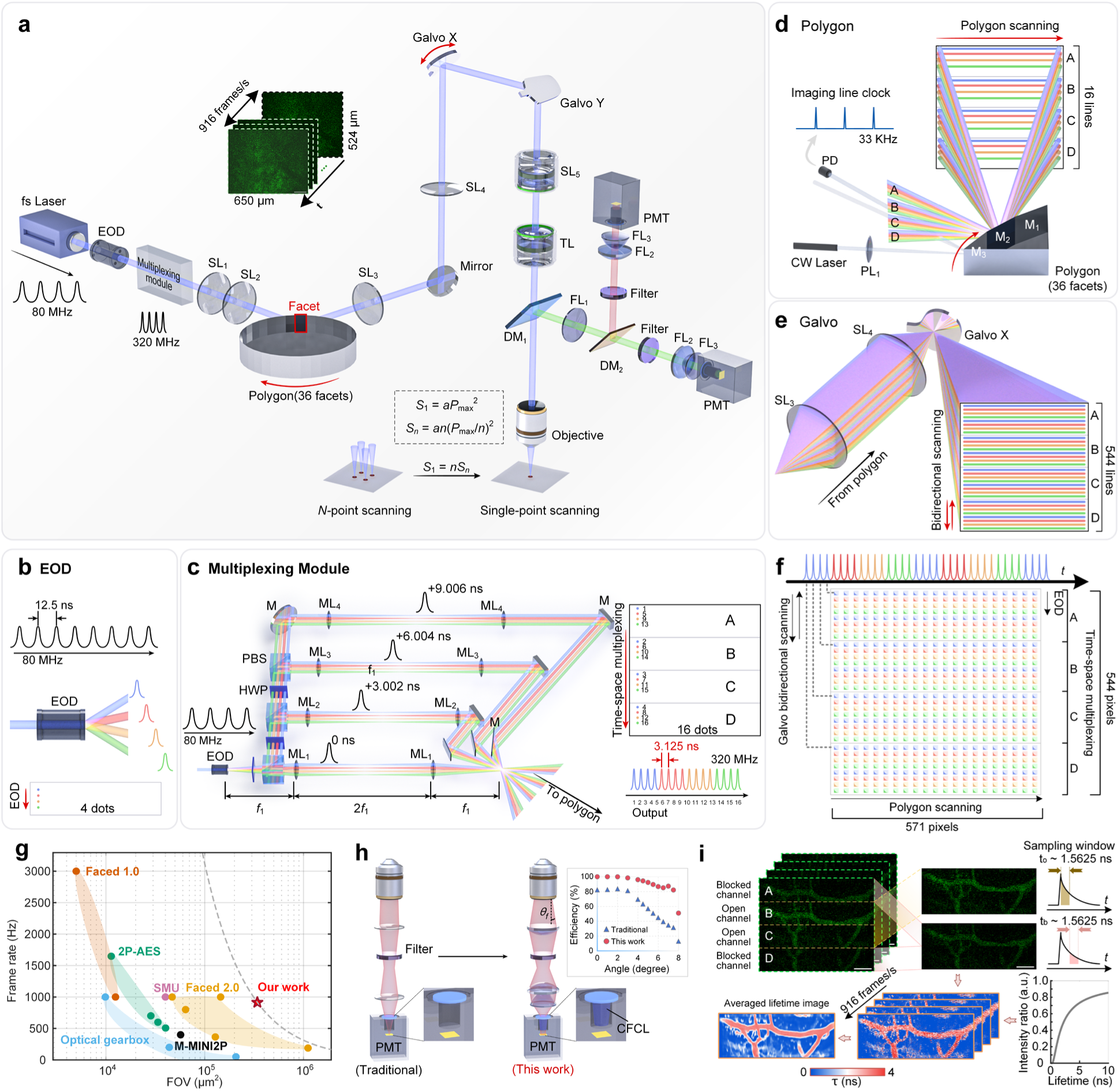
Hybrid scanning architecture and optical design of HS2PM. **a**, Optical layout of the HS2PM system. An 80 MHz femtosecond (fs) laser (920 nm) beam is first angularly encoded by an EOD. It is then split into four time-delayed replicas by the STDM module and scanned along the *x*- and *y*-axes by a polygon and galvo scanner, respectively. This enables 916 Hz raster imaging over a 650 × 524 μm² FOV, with fluorescence collected via a dual-channel wide-angle detection system. **b**, Angular encoding by the EOD generates four spatially offset pulses from the 80 MHz pulse train. **c**, Polarization beam splitters (PBSs), half-wave plates (HWPs), and achromatic delay lines in the STDM module replicate each pulse into four delayed beams (∼3 ns spacing). **d**, The polygon scanner scans all 16 beams to produce 16 lines per facet. A continuous-wave (CW) laser and photodiode (PD) generate synchronization clocks for scan alignment. **e**, Bidirectional galvo scanning fills the *y*-axis gaps between four-line clusters, producing 544 lines per frame. **f**, Excitation pattern per frame, covering the full FOV. **g**, Comparison with other single-point scanning TPVI systems shows HS2PM achieves the largest FOV at near-kilohertz frame rates. **h**, Comparison of emission collection strategies. The custom-designed collection system achieves higher collection efficiency than conventional epi-fluorescence relays within ± 8°. **i**, FLIM mode. STDM channels are reconfigured to generate two temporally offset intensity windows per pixel. Lifetime is estimated from their intensity ratio using a pre-calibrated lookup table.

HS2PM substantially improves photon detection efficiency to boost SNR and depth penetration. Conventional epi-fluorescence relays, typically comprising one or two lenses^7,11,28^, capture only a small fraction of the multiply scattered emission emerging from the objective back aperture at large angles. In HS2PM, we replaced this relay with a three-lens condenser coupled to a custom frustum-of-cone light guide (CFCL), which improves the photon collection efficiency within ± 8° and achieves an overall detection efficiency >80% (Fig. 1h; Extended Data Fig. 3). This configuration substantially increases the number of scattered photons reaching the detector, thereby improving SNR and extending the usable imaging depth at a given excitation power.

HS2PM can also be reconfigured for high-speed fluorescence lifetime imaging (FLIM), enabling *in vivo* estimation of indicator fluorescence lifetimes. In this mode, the STDM output is configured to two active channels, and for each pixel we acquire two temporally separated sampling windows: one corresponding to the prompt emission and the other delayed by approximately 1.56 ns (Fig. 1i; Extended Data Fig. 4b). The fluorescence lifetime at each pixel is then derived from the ratio of the two intensity values, based on the exponential decay kinetics of the fluorophore. Although disabling two STDM channels reduces spatial throughput and halves the accessible FOV, this FLIM mode enables quantitative *in vivo* estimation of indicator fluorescence lifetimes, broadening HS2PM’s functionality beyond intensity imaging.

HS2PM integrates a streamlined data-processing pipeline that preserves millisecond dynamics and enhances neuron identifiability at scale (Extended Data Fig. 5). Raw frames are first rearranged to restore raster images using a pixel-wise order map (Extended Data Fig. 2), with an additional correction for odd-even offsets from bidirectional galvo scanning. STDM crosstalk is then demixed to suppress signal contamination when fluorescence lifetimes exceed the sampling interval (see Methods). Finally, VolPy^29^ is used for motion correction, denoising and robust spike extraction. This end-to-end pipeline ensures high-fidelity voltage readout, enabling reliable population-scale imaging with millisecond precision.

### Stable and deep-layer population voltage imaging with minimal phototoxicity

HS2PM stably resolves millisecond-scale voltage dynamics from large populations of cortical neurons *in vivo*, overcoming the long-standing difficulty of maintaining high-SNR recordings hundreds of micrometers below the cortical surface^6^. Using a soma-targeted voltage indicator (ASAP5-Kv^12^) in awake, head-fixed mice, we imaged a 650 × 524 µm^2^ FOV with a 16× 0.8 NA objective, identifying 165 neurons in a single plane at 490 µm depth under 200 mW excitation (Fig. 2a). Voltage traces from these neurons exhibited robust action potentials with peak -Δ*F*/*F* amplitudes of ∼35% on average (Fig. 2b). Low-pass filtering revealed subthreshold fluctuations, including slow-wave dynamics distinct from suprathreshold spikes. At the population level, spike raster plots revealed coordinated activity (Fig. 2c, top), and the population-averaged firing rate exhibited rhythmic fluctuations (Fig. 2c, bottom). We analyzed population activities from one of our experimental animals, which exhibited more distinct synchronized spiking that was tightly phase-locked to the oscillation cycles (Extended Data Fig. 6). Similar membrane oscillations have previously been observed from neurons in sensory cortex^30^. Single-neuron traces exhibited stereotyped action potential waveforms, with a rapid depolarization followed by a clear after-hyperpolarization, consistent with refractory periods (Fig. 2d). Autocorrelogram (ACG) analysis across the recorded neurons yielded inter-spike interval distributions of approximately ± 2.2 ms (Fig. 2e), confirming the millisecond precision of HS2PM recordings *in vivo*.

**Fig. 2.**
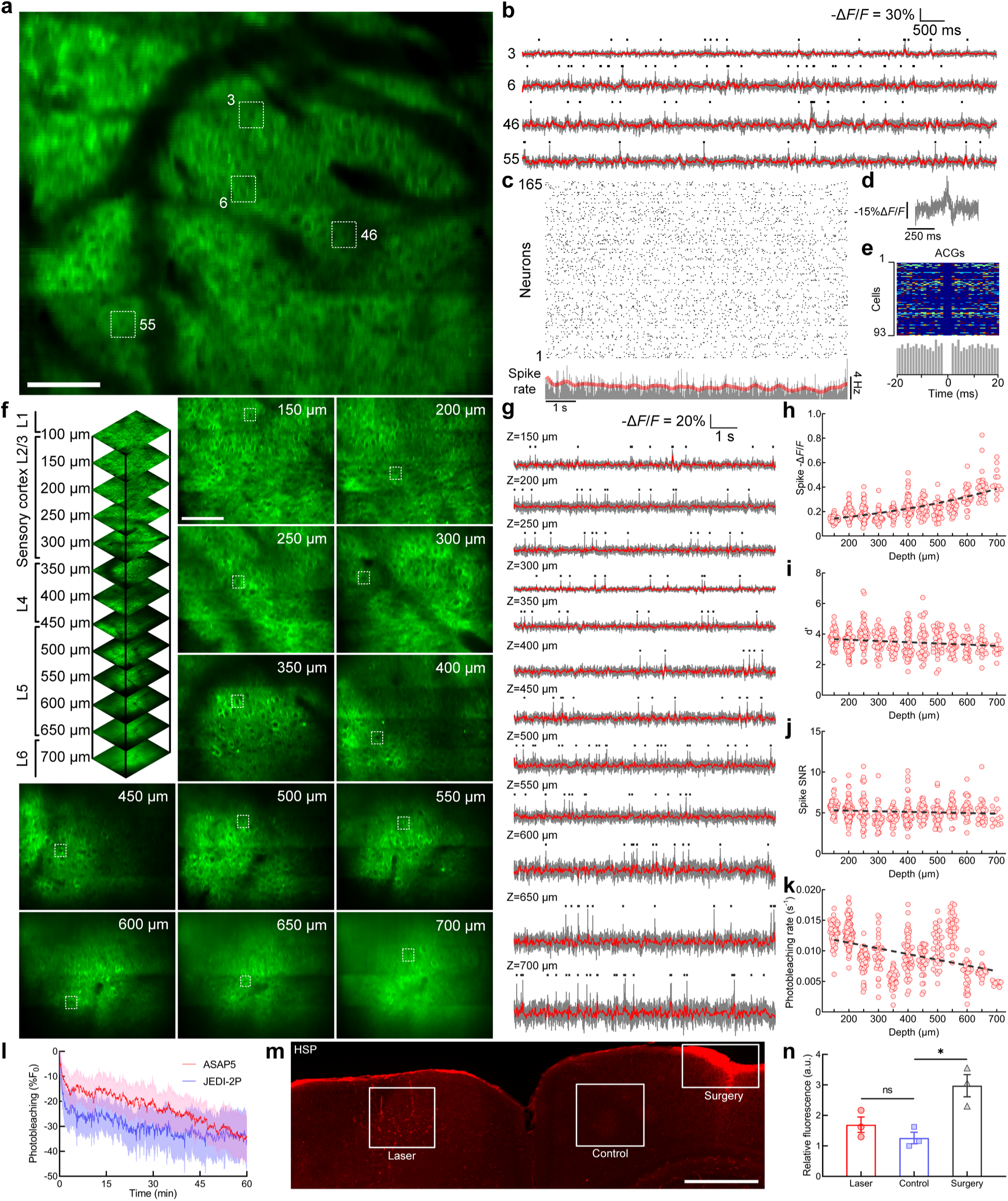
HS2PM enables stable, deep-layer, and long-duration population voltage imaging in awake mouse cortex. **a**, Time-averaged projection of an *in vivo* ASAP5-Kv-labeled image recorded at 490 μm depth in sensory cortex. FOV: 650 × 524 μm^2^ (571 × 544 pixels), acquired at 916 Hz using 920 nm excitation at 200 mW post-objective (Nikon 16×, 0.8 NA). Scale bar, 100 μm. **b**, Representative -Δ*F*/*F* voltage traces (gray) from indexed neurons in **a**, showing robust spike events (black ticks) and subthreshold fluctuations (red). **c**, Raster plot of detected spikes from 165 simultaneously imaged neurons, aligned with the average population firing rate (gray) and its low-pass filtered trace (red). **d**, Example of an individual spike waveform captured in raw fluorescence, exhibiting high temporal resolution and clear peak structure. **e**, ACGs of neurons with ≥20 spikes (each row), indicating the ± 2.2 ms ISI. Bottom histogram: population-averaged ACG. **f**, Time-averaged projection of ASAP5-Kv-labeled neurons across cortical depths (150-700 μm) captured in a 373 × 300 μm^2^ FOV using an Olympus 25×, 1.05 NA objective. Image size: 571 × 544 pixels. Scale bar, 100 μm. **g**, Representative -Δ*F*/*F* voltage traces (gray) and subthreshold components (red) from neurons at various depths in **f**. **h**-**k**, Quantification of spike -Δ*F*/*F*, *d*′, spike SNR and bleaching rate. **l**, One-hour continuous imaging using ASAP5 (*n* = 61 neurons, 2 FOVs, 2 animals) and JEDI-2P (*n* = 55 neurons, 2 FOVs, 2 animals) shows stable performance with < 35% bleaching under 200 mW excitation. Shaded region, s.d. **m**, Immunostaining of HSP-70/72 to assess photodamage. Three cortical regions (white box)—the imaged area, contralateral control region, and surgical injury site—show no significant damage in the imaged area. Scale bar, 500 μm. **n**, Quantification of relative HSP-70/72 expression in imaged and injured regions, normalized to contralateral control (*n* = 3 animals; Welch’s *t* test; \**p* < 0.05, ns = not significant). Error bars: s.e.m.

HS2PM also maintains high-fidelity voltage recordings across cortical layers, achieving reliable spike detection down to 700 µm depth. With a higher NA objective (25×, 1.05 NA; FOV 373 × 300 µm^2^), we captured population-level voltage dynamics spanning layers 1-5 and part of layer 6 (Fig. 2f). Voltage traces from neurons at all depths exhibited robust action potentials (Fig. 2g). Spike -Δ*F*/*F* amplitudes remained consistently high across depths (≥22% on average; Fig. 2h) and showed a slight increase with depth. Spike detection was highly reliable across depths. Although *d*′ declined modestly and quasi-linearly from 150 µm to 700 µm, the mean value remained ∼3.5, and at 700 µm it was still >3 (Fig. 2i), corresponding to a high probability of detection^31^. Spike SNR was largely preserved (∼5) across cortical layers (Fig. 2j), and bleaching rates remained low <0.012 s⁻¹ (Fig. 2k). These results confirm that HS2PM preserves reliable spike detection and signal fidelity in deep cortical layers.

Finally, HS2PM enables long-duration population imaging with minimal bleaching and negligible phototoxicity, overcoming a critical barrier to extended *in vivo* voltage recordings. Continuous 1-h recordings at 200 mW in six mice (three expressing ASAP5-Kv and three JEDI-2P^13^) produced only a modest ∼35% fluorescence decline (Fig. 2l), indicating stable performance with limited bleaching. To further assess phototoxicity, we examined HSP-70/72 expression. Immunostaining revealed no detectable increase in the imaged regions compared to contralateral controls (*n* = 3, *t* = 1.385, *p* = 0.2440, Welch’s t test; Fig. 2m, 2n), whereas strong staining was evident at surgically injured sites caused by head restraint (*n* = 3, *t* = 4.194, *p* = 0.0248). Together, these results establish HS2PM as a robust platform for high-fidelity, long-duration voltage imaging *in vivo*, combining stable population recordings with minimal bleaching and negligible phototoxicity.

### Resolving spikes and subthreshold dynamics during sensory processing

Large-scale calcium imaging has revealed transient reorganizations of cortical population activity during sensory responses^32^. Yet the slow kinetics of calcium indicators obscure the millisecond voltage transients that drive these dynamics, leaving the relative roles of spikes and subthreshold fluctuations unresolved. Using HS2PM, we directly imaged *in vivo* voltage responses to sensory stimulation, capturing both suprathreshold spiking and subthreshold membrane dynamics with millisecond precision.

HS2PM revealed that sustained sensory stimulation drives a rapid and coordinated spiking response across cortical populations. We applied a 3-s continuous air puff (Fig. 3a, top) to head-fixed mice and imaged neuronal voltage dynamics in sensory cortex with a 25× 1.05 NA objective at 355 µm depth (Fig. 3b; 200 mW). Analysis of 9-s voltage traces showed that responsive neurons generated nearly synchronous spikes at stimulus onset (Fig. 3c). This activation was reflected in normalized spike-rate heat maps from 297 neurons in two mice (Fig. 3d), where population-averaged firing rates rose abruptly to ∼9.5 Hz and peaked ∼62.6 ms post-stimulus before relaxing to baseline. Response-latency mapping (Fig. 3e) revealed substantial cell-to-cell variability, yet all activations occurred within 223 ms. Across the entire population, latency histograms exhibited a unimodal peak at 115.6 ms (Fig. 3f), consistent with rapid neuronal adaptation and feed-forward drive^33^. Statistical analyses confirmed that, compared to the period without air puff, the air-puff period showed both an overall increase in spike rate (Fig. 3g) and a modest rise in pairwise correlations (Fig. 3h). Finally, three-dimensional principal component analysis (PCA) trajectories showed a clear transition from pre- to post-stimulus states (Fig. 3i), indicating a pronounced reorganization of cortical activity.

**Fig. 3.**
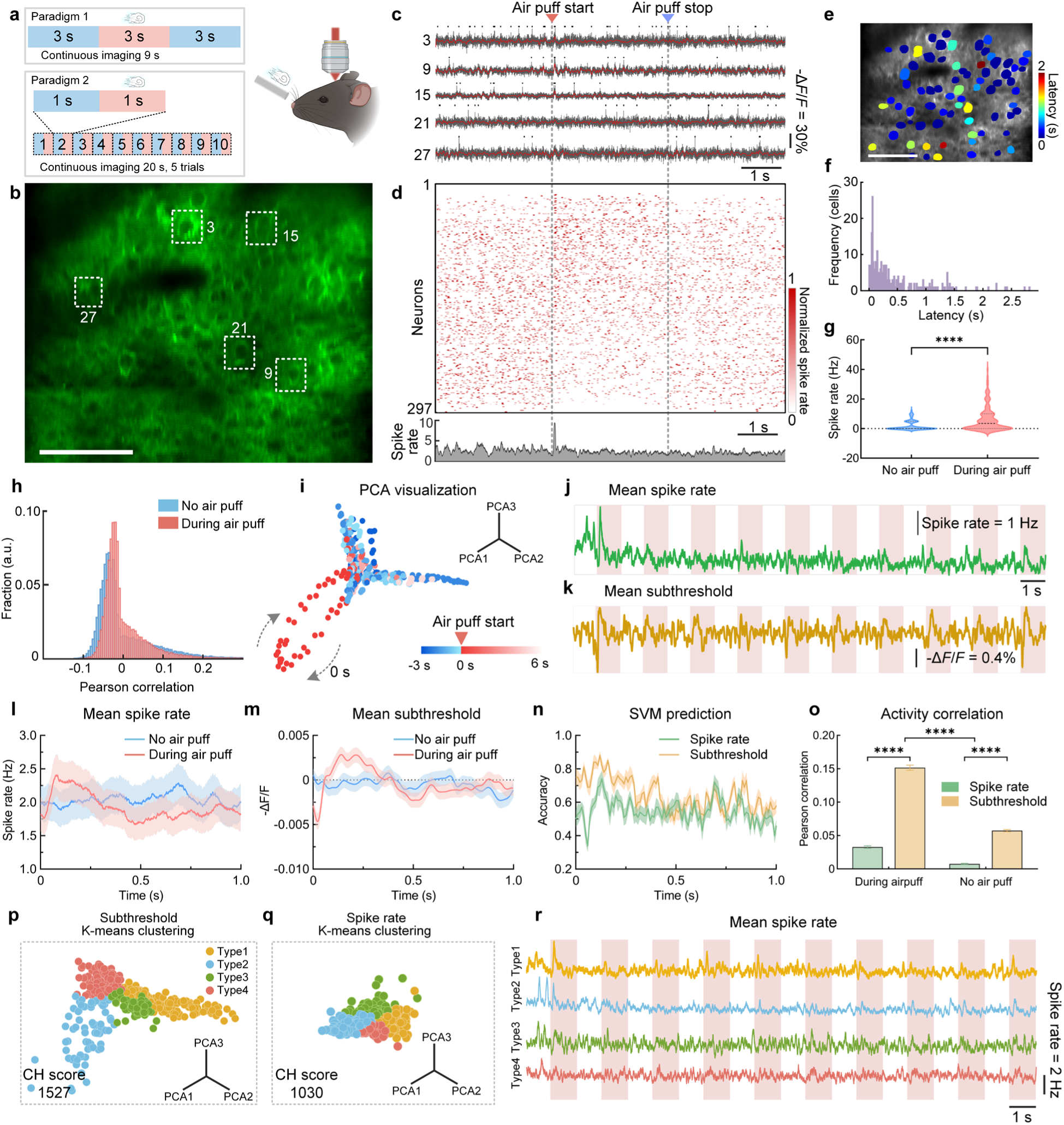
HS2PM resolves rapid cortical voltage dynamics during sensory processing. **a**, Schematic of the two air-puff stimulation paradigms. **b**, Time-averaged image of neurons expressing ASAP5 in sensory cortex (355 μm depth) acquired at 916 Hz (25×, 1.05 NA). **c**, Representative voltage traces from indexed neurons in **b**, showing spikes (black ticks) and subthreshold signals (red) aligned to stimulus onset. **d**, Normalized spike rates of 297 neurons from 2 mice reveal rapid, synchronized responses following stimulus onset; population average shown below. **e**, Latency map of responsive neurons color-coded by time to first evoked spike; see also **f**, histogram of response latencies (peak at ∼116 ms). **g**, Spike rates significantly increased post-stimulation (\*\*\*\**p* < 0.0001, paired t-test). **h**, Pairwise Pearson’s correlation coefficients of spike trains also increased after stimulation. **i**, PCA of population spiking activity revealed a clear trajectory shift following stimulus onset. **j-k**, Repeated stimuli evoked robust initial spiking (**j**) but stable subthreshold responses across trials (**k**). **l-m**, Both spikes and subthreshold signals increased during stimulation and returned to baseline. **n**, Classifiers trained on subthreshold signals outperformed those based on spike rates in decoding stimulus identity (SVM accuracy: 64% vs. 55%). **o**, Air puff increased neuron-population synchrony in both spike rate and subthreshold domains (\*\*\*\**p* < 0.0001, 2-way ANOVA). **p-q**, K-means clustering based on subthreshold dynamics yielded more cohesive and interpretable response types (Calinski-Harabasz score: 1527 vs. 1030). **r**, Subthreshold-based clusters exhibited distinct adaptation patterns across repeated stimuli: Type 1 neurons remained responsive, while Types 2/3 adapted and Type 4 was non-responsive. Scale bars: 100 μm in **b** and **e**. Shaded region in **l**-**n**, s.e.m.

Repeated stimulation revealed a dissociation between spiking adaptation and stable subthreshold dynamics. We applied a 10 × 1-s air puffs within 20 s (Fig. 3a, bottom), across five trials per FOV. Each puff evoked an immediate spike-rate increase, with the strongest response during the first puff and progressively attenuated activity thereafter (Fig. 3j). In contrast, subthreshold voltage traces exhibited reproducible -Δ*F*/*F* transients for every puff, with amplitudes remaining stable across repetitions (Fig. 3k). At the population level, both spike rates and subthreshold signals rose physically at puff onset and decayed back to baseline (Fig. 3l and 3m), confirming consistent stimulus encoding across suprathreshold and subthreshold domains. Quantitative analyses further highlighted the informational advantage of subthreshold signals: support vector machine (SVM) classifiers trained on subthreshold traces outperformed those based on spike-rate vectors (64.01 ± 4.42 % vs. 54.61 ± 4.54 %, Fig. 3n), and both spike and subthreshold correlations with the surrounding population increased significantly during stimulation (r_rate_ = 0.0327 ± 0.0019 vs. 0.0074 ± 0.0008; r_sub_ = 0.1515 ± 0.0038 vs. 0.0573 ± 0.0016; *p* < 0.0001; Fig. 3o). Together, these results demonstrate that while spikes adapt under repeated stimulation, subthreshold dynamics remain robust, supporting enhanced network-level coordination.

Subthreshold activity provided a more reliable basis for classifying functional response types. We applied unsupervised k-means clustering to neurons grouped by either spike-rate profiles or subthreshold voltage dynamics. Both approaches partitioned the population into four functional classes. Subthreshold-based clustering produced higher intra-cluster cohesion, as quantified by the Calinski-Harabasz (CH) score (1527 vs. 1030; Fig. 3p,q), and produced distributions that were more consistent with biologically interpretable response types (Type 1-4: *n*_1_ = 210, *n*_2_ = 82, *n*_3_ = 60, *n*_4_ = 233 in Fig. 3p vs. *n*_1_ = 158, *n*_2_ = 254, *n*_3_ = 97, *n*_4_ = 76 in Fig. 3q). We therefore adopted the subthreshold classification for subsequent analyses. The four classes exhibited distinct response patterns to repeated air puff stimuli (Fig. 3r): Type 1 neurons responded robustly to every puff, Type 4 neurons remained largely inactive, and Types 2 and 3 showed strong initial responses that rapidly attenuated with repetition. These results demonstrate that subthreshold-based clustering yielded higher fidelity and revealed functional heterogeneity in neuronal adaptation to repeated stimuli.

Together, HS2PM resolves millisecond-scale reorganization of cortical activity and reveals how spikes and subthreshold dynamics jointly contribute to population coding, providing insights into sensory processing that remain inaccessible with calcium imaging.

### Population-scale vascular imaging and high-speed fluorescence lifetime imaging

HS2PM enables population-scale vascular imaging by overcoming the one-vessel limitation of conventional two-photon line-scanning approaches^34,35^. Whereas traditional methods restrict flow measurements to a single vessel, the near-kHz frame rates of HS2PM provide the temporal resolution needed for high-throughput analysis of flow velocity and diameter across many vessels simultaneously. To demonstrate this capability, we labeled the cerebral vasculature in mice with Rhodamine-B and recorded vascular fluorescence at 916 Hz, using a 25× objective at 90 µm depth (100 mW; Fig. 4a). Millisecond-resolved traces from two representative vessel segments revealed flow fluctuations with average velocities of 0.0875 mm/s and 0.0935 mm/s, respectively (Fig. 4b).

**Fig. 4.**
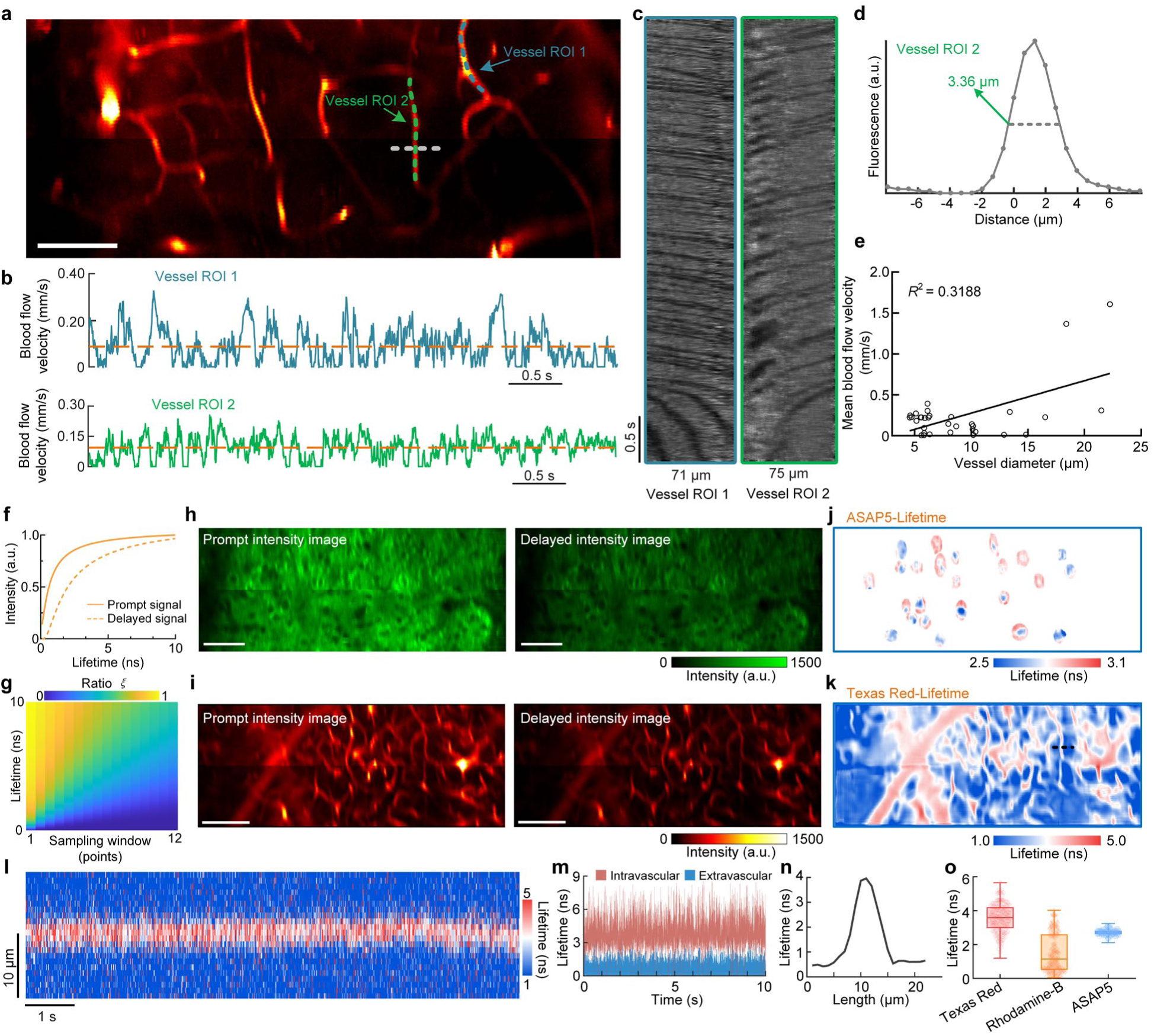
HS2PM enables high-speed population-scale vascular imaging and pixel-resolved fluorescence lifetime mapping. **a**, Time-averaged projection of vasculature labeled with Rhodamine-B at 100 μm depth. Two vessel segments (ROIs) are traced for flow analysis. **b**, Flow velocity fluctuations over 5 s from the two vessel ROIs. **c**, Kymographs showing RBC movement as diagonal streaks. **d**, Vessel diameter extracted by Radon transform and FWHM. **e**, Positive correlation between vessel diameter and flow velocity (*n* = 33 vessels). **f-g**, FLIM calibration curves and color-coded map linking fluorescence lifetime to intensity ratio (*ξ*) across dual time-offset channels (see methods for details). **h-i**, Prompt and delayed emission frames of ASAP5-expressing neurons (185 μm depth) and Texas Red-labeled vasculature (90 μm depth). **j-k**, Corresponding pixel-resolved lifetime maps. **l-n**, Lifetime contrast across vessel cross-section reveals elevated lifetime within lumen; fluctuations and spatial profile are consistent with erythrocyte flux. **o**, Summary of lifetime measurements across indicators: Texas Red, Rhodamine-B, and ASAP5 (*n* = 86, 86, 58 ROIs). Scale bars: 50 μm in **a** and **h**, 100 μm in **i**.

Spatiotemporal kymographs visualized red blood cell movement as diagonal streaks (Fig. 4c), while vessel diameter was extracted from fluorescence cross-sections using Radon transform and defined as the full-width at half-maximum (FWHM; Fig. 4d). Across 33 vessel segments, HS2PM enabled simultaneous quantification of flow velocity and vessel diameter, revealing a positive correlation between the two metrics (*R*^2^ = 0.3188; Fig. 4e), consistent with physiological expectations. These results establish the feasibility of population-scale vascular flow measurements with HS2PM.

HS2PM extends beyond intensity imaging by enabling high-speed, pixel-resolved FLIM. We implemented a dual-channel ratio method in which fluorescence was acquired in two temporally offset sampling windows, corresponding to prompt and delayed emission (Extended Data Fig. 4b). The ratio *ξ* between these signals directly encodes fluorescence lifetime based on exponential decay kinetics (Fig. 4f), thereby converting conventional intensity images into quantitative, pixel-resolved lifetime maps (Fig. 4g). Applying this approach, we resolved lifetime contrasts in both neurons and vasculature. ASAP5-labeled neurons and Texas Red-labeled vessels produced distinct peak and faded intensity images (Figs. 4h,i), which were converted into pixel-resolved lifetime maps (Figs. 4j,k). In vessel cross-section, lifetime mapping revealed clear contrasts between intravascular and extravascular regions: the vessel lumen exhibited fluctuating lifetimes, whereas surrounding tissue showed consistently short values (∼0.58 ns; Fig. 4l,m,n), consistent with endogenous autofluorescence. Finally, quantitative analysis across regions of interests (ROIs) confirmed robust fluorophore-specific lifetimes: Texas Red- and Rhodamine-B-labeled vasculature exhibited mean values of 3.57 ± 0.07 ns and 1.50 ± 0.10 ns, respectively, while ASAP5-expressing neurons showed 2.72 ± 0.01 ns (mean ± s.e.m.; Fig. 4o), consistent with prior benchmarks^36–38^.

## Discussion

We present HS2PM, a hybrid scanning two-photon microscope that resolves the long-standing trade-off between imaging speed, FOV, and excitation efficiency in TPVI. By preserving the photon efficiency inherent to single-point excitation while achieving kilohertz frame rates across a population-scale FOV, HS2PM enables reliable *in vivo* recording of both spiking and subthreshold voltage dynamics with single-cell and one-millisecond resolution.

This study introduces a conceptual and technical framework that redefines the design constraints of TPVI. Through theoretical modeling, we demonstrate that the fundamental limitation lies not only in scanning speed but also in photon flux under the biological power ceiling. Our analysis shows that within realistic power limits, single-point excitation offers the most practical route to achieving sufficient excitation efficiency for reliable voltage detection (Supplementary Note 2). This underscores that excitation efficiency is as critical as frame rate. Based on this insight, we developed a hybrid scanning strategy that overcomes the intrinsic speed limit of point scanning. Operating near the theoretical upper bound imposed by fluorophore lifetimes of approximately 3 nanoseconds, HS2PM achieves kilohertz-rate imaging at 320 MHz effective sampling without compromising FOV, thereby achieving the highest sampling rate and the largest FOV among existing single-point scanning methods^11,20,25–27^. The resulting square-format population-scale imaging field allows seamless adaptation of established calcium imaging paradigms to voltage imaging. In addition, the same temporal multiplexing framework supports high-speed fluorescence lifetime imaging, enabling kilohertz-rate lifetime mapping without requiring electro-optic gating^39^ while also integrating deep, fast, and quantitative voltage imaging within a single system.

By scaling voltage imaging to the population level, HS2PM enables circuit-level analyses that were previously limited to calcium imaging, but with millisecond precision sufficient to resolve both action potentials and subthreshold membrane fluctuations *in vivo*. This capability allows direct dissection of how spikes and subthreshold dynamics jointly shape sensory adaptation, network coordination, and population coding. These results provide new experimental access to fundamental questions in systems neuroscience, for example how neural circuits compute^40^, how subthreshold activity contributes to collective dynamics^41,42^, and how excitatory and inhibitory inputs interact to shape cortical responses^42^, questions that have remained inaccessible with calcium imaging alone.

Several limitations remain. The fluorescence lifetime of existing GEVIs (∼3 ns)^38^ restricts further gains in temporal multiplexing and introduces signal crosstalk^43^, necessitating demixing algorithms for correction. In addition, the 36-facet polygon scanner imposes a physical constraint on FOV, though recent two-pass geometries suggest this could be expanded up to twofold^27^. Biologically, variable GEVI expression levels and the absence of demonstrations in chronic or freely moving models remain critical challenges to broader application.

Future directions include pairing HS2PM with next-generation voltage indicators that offer shorter lifetimes, enhanced brightness, or red-shifted spectra. This would further improve imaging speed, depth, and multiplexing capabilities. Beyond neuronal populations, HS2PM may be applied to vascular dynamics, neuron-glia interactions, and all-optical physiology. The system provides a flexible platform for multimodal interrogation of neural circuits. With continued development, HS2PM has the potential to achieve population-scale millisecond-resolved imaging of brain activity in both healthy and diseased states. This positions voltage imaging to fulfill a transformative role in neuroscience research comparable to that of two-photon calcium imaging over the past two decades.

## Methods

### HS2PM microscope

We simulated and optimized the placement of optical components using Zemax OpticStudio software (Zemax LLC). The complete design was then assembled in a 3D SolidWorks environment to verify there was no component interference. Custom aluminium holders and frames were machined for rigid mounting of the optics. A detailed schematic of the HS2PM setup is shown in Extended Data Fig. 1 (see the animated rendering in Supplementary Video 1), with components listed in Supplementary Table 2. A 920-nm, 2-W fiber laser (ALCOR920, Sparks Lasers; 100-fs pulse duration, 80-MHz repetition rate) was used. The laser spot was focused onto the sample via optical scanning devices, relay lenses and an objective (CFI75 LWD 16X W, Nikon; 16×/0.8 NA). The 1/e^2^ beam diameter at the back aperture is ∼5.31 mm, leading to an excitation NA of ∼0.28. A 1.05 NA objective (XLPLN 25xWMP2, Olympus; 25×) was used for higher-resolution voltage imaging with an excitation NA of ∼0.46. The post-objective power was controlled by the acousto-optic modulator (AOM) integrated in the laser.

The EOD (Model 400-120, Conoptics; 8.5 μrad/V deflection efficiency) was driven by an amplifier (Model 25A, Conoptics) synchronized to the laser clock. An arbitrary waveform generator (AWG; including PCI-8361, PXIe-8360 and PXIe-1062Q; National Instruments) output 20 MHz periodic waveforms, switching the voltage applied to the EOD every 12.5 ns. By cycling through four distinct voltage levels, the EOD deflected successive laser pulses into four longitudinal positions. These beams then entered the multiplexing module, which further split each into four spatially separated channels via a combination of time- and space-division multiplexing. This produced four vertical excitation points per scanning region, totaling 16 aligned points across the full FOV.

The multiplexing module consisted of three half-wave plates (HWP; 2 × WPH05M-915, 1 × WPH10M-915, Thorlab), three polarizing beam splitter cubes (PBS; PBS105, Thorlabs), eight lenses, and several mirrors. Three HWPs were individually placed before each PBS to balance optical power across the four output paths. The STDM module employed four sets of achromatic doublets (listed in Supplementary Table 2), with each set arranged in a 4F configuration and optimized with channel-specific focal lengths. Differential path lengths of 900 mm between adjacent channels introduced ∼3 ns inter-pulse delays, raising the effective pulse repetition rate to 320 MHz. This value was chosen to approach the theoretical limit for single-point excitation, which is set by the fluorescence lifetime of the indicators at a ∼3 ns inter-pulse interval^38^, and to minimize fluorescence crosstalk. To minimize aberrations across the FOV, the EOD and multiplexing module were mounted on an inclined metal plate tilted at ∼1.465° (Extended Data Fig. 1d), ensuring symmetric beam arrangement relative to the subsequent optical axis. Due to small inter-channel angles and proximity to the beam convergence point, the second lens in the first channel was trimmed into a 3.5-mm-wide strip to avoid obstructing adjacent beams (Extended Data Fig. 1e).

The output beam from the multiplexing module was first reduced in diameter by a two-lens group and reimaged onto a 36-facet polygon scanner (SA34, Lincoln Laser) rotating at 54,945 rpm (916 Hz). The imaging frame rate matched the polygon rotation frequency. The beam was then relayed to a galvanometer scanner pair (6215H, Cambridge Technology). The Galvo-X scanner performed *y*-axis scanning, with its deflection angle set by inter-channel angular differences. To improve speed and stability, Galvo-X scanner operated in a vertical bidirectional scanning mode at half the frame rate (top-down for odd frames, bottom-up for even frames), minimizing flyback dead time (see Supplementary Fig. 1). Galvo-Y scanner remained stationary during voltage imaging due to its slower response. The last two facets of each polygon rotation were allocated to accommodate Galvo-X flyback, resulting in 34 effective scanning facets per frame. It was multiplied by 16 through the combined action of the EOD and STDM, yielding a total of 544 scan lines covering a 524 μm vertical FOV per frame.

To reduce system cost, two identical TTL200MP lenses (Thorlabs) were used as the scan and tube lenses. A dichroic mirror (DM_1_; FF735-Di02-25x36, Semrock) transmitted excitation light to the objective while reflecting fluorescence toward the detection path. A second dichroic mirror (DM_2_; FF552-Di02-25x36, Semrock) then split the fluorescence into two spectral channels. Each channel was equipped with a photomultiplier tube (PMT; H7422-40, Hamamatsu) coupled to an amplifier (C11184, Hamamatsu; DC-300 MHz), and signals were digitized at 2.56 GHz using a vDAQ™-HS system (Vidrio Technologies).

In addition to kilohertz voltage imaging, the HS2PM system supported low-frame-rate and bright-field imaging modes. For low-frame-rate scanning, the system switched to a conventional two-photon mode: the polygon scanner and EOD were disabled, and only one channel in the multiplexing module remained open, while the Galvo-X and Galvo-Y scanners worked in tandem. This configuration provided a large FOV of 2.06 × 2.06 mm^2^ (1024×1024 pixels, 1.71 Hz, Supplementary Fig. 2a), enabling rapid target identification after which the system could switch back to kHz voltage imaging of the region of interest (see Supplementary Video 2 for comparison). The system also offered a bright-field imaging mode, where an LED source was used. Under remote control, the movable mirror (MM; Extended Data Fig. 1c) could be moved into the path between the scan and tube lenses, blocking the 920 nm laser and directing LED light to the objective. Simultaneously, DM_1_ was removed from the optical path, allowing sample-reflected light to reach the eyepiece or CCD camera for bright-field visualization.

### Optimized fluorescence collection system

The dichroic mirror split the collected fluorescence into two channels, each equipped with a bandpass filter (FF01-593/46, FF01-520/35-25, Semrock). Both channels featured identical optical assemblies. Fluorescence exiting the objective’s back aperture first passed through a 75 mm focal length plano-convex lens (LA1386-A, Thorlabs; Ø38.1 mm). The beam then traversed a second plano-convex lens (LA1274-A, Thorlabs; *f* = 40 mm, Ø30 mm) and an aspheric condenser lens (ACL25416U-A, Thorlabs; *f* = 16 mm, Ø25.4 mm), spaced 1 mm apart. This dual-lens configuration coupled the fluorescence to a custom conical frustum light guide (front surface Ø5.85 mm, back surface Ø4.45 mm). A critical innovation involved integrating the CFCL into the 7.4 mm pathway between the PMT’s front window and its internal protective window. For assembly, the 8.5 mm long CFCL was coaxially bonded to a glass disc (Ø14 mm, 1 mm thickness). An annular spacer was first placed at the original PMT window position to create a buffer zone. The CFCL-disc assembly was then inserted into the pathway, with UV-curable adhesive used to co-axially bond the PMT housing, spacer, and glass disc.

We performed Zemax non-sequential ray-tracing simulations of fluorescence emission at varying angles from the objective’s back aperture. The simulations utilized an Embedded Monte Carlo approach, tracing 5×10^5^ rays per angle at a wavelength of 550 nm. Collection efficiency at each emission angle was determined as the ratio of optical power recorded by the detector to the 1-W input power (Extended Data Fig. 3).

### Fluorescence crosstalk characterization

To quantify temporal and spatial crosstalk in scattering tissue, we imaged a cranial-window-implanted mouse expressing ASAP5. For each measurement, only one scanning region was excited at ∼50 mW by enabling a single channel of the multiplexing module, while fluorescence was collected from all four channels. This procedure was repeated sequentially for each of the four channels. At each imaging depth from 150 to 700 μm below the pial surface (in 50 μm steps), 10,000 frames were acquired at 916 Hz, and average intensity images were generated (see Extended Data Fig. 7a). It should be noted that while the schematics in Fig. 1 and Extended Data Fig. 2 illustrate four scanning regions (A-D) for conceptual clarity, the actual spatiotemporal arrangement of the four channels after the multiplexing module is shown in Extended Data Fig. 7a, where channels 1-4 represent the true scanning sequence and spatial layout. Pixels within the excited subregion corresponding to ASAP5 fluorescence were defined as those whose intensities ranked among the top 5%. To quantify crosstalk, the mean intensity of these bright pixels in the excited subregion was compared to the mean intensity of spatially corresponding pixels in each non-excited subregion. The crosstalk percentage was calculated as the ratio of the non-excited to excited mean intensities (see Extended Data Fig. 7b).

### Synchronization and data acquisition

A complete synchronization diagram of the scanning and acquisition system is provided in Extended Data Fig. 8. Polygon scanner synchronization was achieved using a 780-nm reference laser (LM15-780, Lbtek) focused via a plano-convex lens (LA1484-A, Thorlabs) onto the polygon mirror (Extended Data Fig. 1f). The reflected beam was detected by a photodiode (S2506-02, Hamamatsu) mounted on a dedicated circuit board (3-1-5248-900-00, Cambridge Technology), which generated a TTL start-of-line (SOL) pulse at each polygon facet transition. This SOL pulse triggered the data acquisition system, ensuring temporal alignment between the scanning trajectory and signal digitization. The laser clock was synchronized with the SOL signal, galvanometer drivers, AWG, and acquisition card to generate raw data frames that were subsequently reconstructed into spatially accurate images.

The high-speed acquisition system could sample each laser pulse 32 times, but the data acquisition setting was configured differently for voltage imaging and FLIM modes. In voltage imaging, acquisition windows for the four multiplexed channels were strategically aligned to capture each channel’s peak signal. Each window spanned 4 consecutive sampling points within the 32-point pulse cycle, separated by 4-point intervals to minimize inter-channel crosstalk (Extended Data Fig. 4a). This configuration enabled full-FOV imaging at 916 Hz. In FLIM mode, the system was reconfigured to estimate fluorescence lifetimes *in vivo* by blocking two non-adjacent multiplexing channels (e.g., channels 2 and 4), effectively imaging half of the full FOV. The remaining two active channels operated with a 6.25 ns pulse interval. Each pixel was acquired by two sampling windows: one capturing the prompt intensity and the other capturing the intensity after a 1.5625 ns delay (Fig. 1i; Extended Data Fig. 4b). The intensity ratio between these windows was then used for pixel-wise lifetime calculation.

### FLIM

Given that fluorescence emission follows an exponential decay, we use *τ* to represent fluorescence lifetime, and the fluorescence intensity *F*(t) can be expressed as *F*_0_e^-*t*/*τ*^. Fluorescence was simultaneously sampled by two contiguous, width-matched detection windows, producing two subareas’ images corresponding to the prompt (P) and delayed (D) intensity:

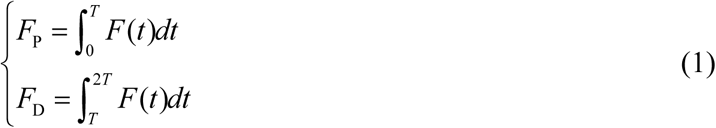

where *T* presents the equal-time ∼1.5625 ns during each sampling window. The intensity ratio *ξ* between the two adjacent sampling windows directly correlates with the fluorescence lifetime:

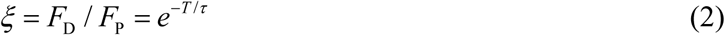

Therefore, the two-subarea pair encoded nanosecond time information in their intensity ratio. The relationship curve between *ξ* and *τ* is shown in Fig.1i. We can compute the lifetime values for each pixel using this intensity ratio, and the principle is similar to existing wide-field lifetime imaging techniques^39^.

### Animal preparation

All mice used were male and of a C57BL/6JNifdc background, purchased from Charles River Co., Ltd. All mice were socially housed in the Animal Facility at the Department of Laboratory Animal Science at Fudan University under a 12 h light/dark cycle (lights on 08:00, off 20:00) and room temperature with food and water ad libitum. All animal experiments were conducted in accordance with the guidelines of the Institutional Animal Care and Use Committee of the Department of Laboratory Animal Science at Fudan University (2022JSITBR-013, 2024-JSYY-035). *In vivo* voltage imaging data in this study were collected from 9 wild-type mice, six eight-week-old male mice were injected with pAAV-CaMKiia-ASAP5-Kv (a generous gift from Michael Z. Lin in Stanford University) in the sensory cortex (2/1.5 mm left medial-lateral and - 1.06 mm anterior-posterior to lambda), and the other three were injected with AAV-CaMKii-JEDI-2P-Kv in the same site at the same age. The microinjection pump (KDS 130, KD Scientific) was used to inject at a rate of 50 nl/min at each viral injection site, with a total volume of 200 nl using a 10 μl Hamilton syringe (Model 701N). For visualization of GEVI-labeled neurons via fluorescence microscopy, the mice underwent cranial window implantation surgery after 2 months for sufficient expression of the indicators. In brief, mice aged 4 months were anesthetized with 1∼2% isoflurane (RWD Life 524 Science #R510-22-10) and head-fixed in a stereotaxic apparatus, then a 4 mm circular craniotomy was made using a high-speed dental drill over the virus injection position. A glass window made of a single coverslip customized by us was flushed with the skull, and sealed by a tissue adhesive (VetBond, 3M). A stainless-steel head-fixator was then firmly attached to the skull with dental acrylic and cyanoacrylate adhesive (ergo, 5800). Implanted mice were allowed a one-week recovery period, during which they received daily intraperitoneal injection of Dexamethasone (ST1254, Beyotime Biotechnology) and Cefazolin (Shanghai Macklin Biochemical Technology #HY-B0712A). Both drugs were prepared as 0.5 mg/ml solutions, and each mouse was injected 0.1 ml of each drug. Two-photon calcium imaging was conducted in a 6-month-old male transgenic CaMKii-Cre::Ai162 mice derived from the crossbreeding of CaMKii-Cre (#C001015, from Cyagen Biosciences) and Ai162 mice (#031562, from Jackson Labs). All surgical procedures and pharmacological manipulations followed the same protocol used for the voltage indicator-labelled mice, except that a 6-mm-diameter craniotomy was made over the central brain region. Mice were then adapted to head fixation for 30 mins per day over one week before imaging. After imaging experiments, unless specified, experimental animals were euthanized by CO₂ asphyxiation followed by cervical dislocation.

### *In vivo* voltage imaging

Voltage imaging was performed on awake, head-fixed mice using a total excitation power of 200 mW (measured post-objective). To characterize the imaging depth of HS2PM, spontaneous neuronal activity was recorded at 50 μm intervals starting from 150 μm below the pial surface until the signal could no longer be clearly resolved. At each depth, 10-second movie was acquired at 916 Hz for subsequent analysis of depth-dependent signal characteristics.

For sensory-evoked voltage recordings, two distinct whisker stimulation paradigms were applied contralateral to the imaged hemisphere. The first paradigm consisted of alternating periods of 3 s baseline, 3 s air-puff stimulation, and another 3 s post-stimulation, forming a single 9-s imaging trial. Each mouse was imaged across three different FOVs, with 10 trials recorded per FOV. The second paradigm featured transient whisker stimulation during 20-s continuous imaging sessions, using repeated 1-s air puffs with 1-s inter-stimulus intervals. For this paradigm, five trials were acquired per FOV across three FOVs per mouse. All stimuli were delivered at ∼30 psi through a 20-gauge blunt needle positioned 1-2 cm from the mouse’s face.

### Image processing and analysis

The HS2PM image processing workflow consists of the following steps: raw data rearrangement, crosstalk removal, motion correction, denoising and extraction of neuronal voltage traces (Extended Data Fig. 5). Each raw data frame initially contained 2284 × 34 pixels. Each raw frame represented one of four multiplexed scanning regions within the full FOV. To reconstruct complete frames, four consecutive raw frames were combined into a 571 × 544 composite image at the system’s native 916 Hz acquisition rate. Since the system employed bidirectional Galvo-X scanning—with opposite scanning directions for odd and even frames—a correction algorithm was applied during frame rearrangement to eliminate jitter artifacts caused by the alternating scan orientations. A custom algorithm was developed to remove crosstalk arising from the STDM scheme. This crosstalk was observed between temporally adjacent channels and negligible between non-adjacent channels. The relationship between the measured signal in each channel (CH*_i_*, *i*=1, 2, 3, 4) and the underlying crosstalk-free signal (P*_j_*, *j*=1, 2, 3, 4) was modeled as a linear mixing process:

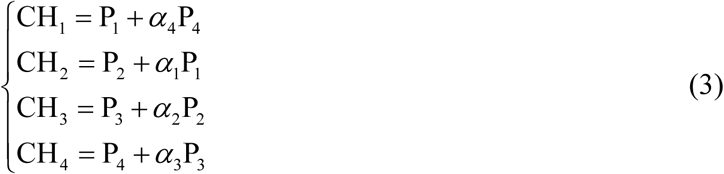

in which *α_i_* is the crosstalk ratio of each temporal adjacent channel. The demixed crosstalk-free frame was obtained by solving the equations, and the images before and after crosstalk demixing process were presented in Extended Data Fig. 5c and 5d for comparison. The original crosstalk ratios were determined by the measured crosstalk in Extended Data Fig. 7, and then optimized during the demixing process. Neurons were identified and selected manually on the averaged images. Finally, the Volpy algorithm was used for motion correction, denoising and extracting neuronal voltage activity traces.

For voltage imaging data analyses, -Δ*F*/*F* was calculated as the negative fractional change in fluorescence relative to the baseline level. The fluorescence baseline was obtained by applying a rolling percentile filter (50%, 500-ms window) to the mean-intensity trace of the ROI. We classified optically identified action potentials (APs) as voltage transients only if they were labeled as spikes by the VolPy algorithm and their peak -Δ*F*/*F* exceeded 2.5 times the standard deviation of the voltage trace within a 20-frame (∼21.84 ms) rolling window centered on the spike. Subthreshold -Δ*F*/*F* was calculated through the application of a 20-Hz, 5th-order Butterworth lowpass filter to the -Δ*F*/*F* voltage trace. Spike rate traces were obtained by applying a rolling 20-ms window on the -Δ*F*/*F* voltage traces and calculating the spike rate within this window. Alongside the -Δ*F*/*F* traces, the SNR traces were calculated as the ratio of Δ*F* (the functional change in fluorescence) to 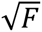 (Poisson noise). The neuronal spike SNR was determined by averaging the SNR values at the time points of identified spikes. Photobleaching rate was quantified as the slope of a linear fit to the normalized fluorescence decay relative to its initial intensity. For sensory-evoked responses, neuronal latency was defined as the interval from stimulus onset to the first subsequent action potential. For calcium indicator-labeled neurons, the baseline fluorescence was calculated as the mean intensity of the initial 30% of each recording trace. All statistical analyses were conducted using MATLAB (R2022b, MathWorks) or built-in tests in SPSS (v20, IBM).

### Photobleaching

Photobleaching measurements were conducted using the HS2PM on male adult mice with implanted cranial windows. The tissue was scanned and imaged at a frame rate of 916 Hz using a total laser power of 200 mW at the sample, and the scanning imaging was performed continuously for 60 min. ROIs were extracted manually. Each data point in Fig. 2l corresponded to the mean fluorescence intensity of every 50th frame across one session of imaging. Photobleaching rates were determined by normalizing all data points to the first data point.

### Photodamage

Three adult mice injected with virus and implanted with cranial windows were used for the experiments. The laser exposure location was selected away from the viral injection sites: we first found labeled neurons by using the microscope and then moved the stage until no neurons were found in the FOV. For each photodamage session, continuous laser scanning was performed for 1 hour at a total power of 200 mW. Twenty hours after laser exposure, mice were anesthetized with 1% pentobarbital sodium and transcardially perfused with heparinized saline (20 U/mL), followed by 4% paraformaldehyde (PFA). Cranial windows were removed to expose the cortex. Brains were postfixed in 4% PFA at 4 °C overnight, then sequentially cryoprotected in 20% and 30% sucrose/PBS solutions, and sectioned into slices with the thickness of 25 µm using a vibratome. To assess photodamage after 1 h of continuous imaging through immunostaining, brain slices were first incubated in blocking solution (5% normal donkey serum/PBST, where PBST was PBS supplemented with 0.3% Triton X-100) for 1 hour. Slices were then incubated with the primary antibody (monoclonal antibody against HSP70/HSP72, Enzo Life Sciences #ADI-SPA-810, diluted at 1:400) at 4 °C overnight, followed by labeling with the secondary antibody (Alexa Fluor® 594 AffiniPure® Donkey Anti-Mouse IgG (H+L), Jackson #715-585-151) for 2 hours at room temperature. Slices were mounted with Antifade Mounting Medium containing DAPI (Beyotime Biotechnology #P0131). Olympus SLIDEVIEW VS200 was used to image the slices with Olympus OlyVIA software. Photodamage was assessed by comparing antibody labeling intensity in the laser-exposed region and its corresponding area in the contralateral hemisphere. Fluorescence intensity was measured in three cortical regions per slice: the laser exposure site, a contralateral control area at least 1 mm away from the laser exposure site (to account for potential effects of chronic window implantation or virus injection unrelated to laser exposure), and the surgically injured region for chronic window implantation. For each region, relative fluorescence was calculated by taking the ratio of its average fluorescent intensity to that of the control area. Data analysis was performed using Welch’s t test (*n* = 3 mice).

### Neural decoding and clustering

To assess how population activity encodes air puff stimuli, we trained linear SVM decoders^44^ on two representations: population spike-rate vectors and subthreshold Δ*F*/*F* traces. Data were collected from 3 mice (9 FOVs, 5 trials per FOV). For each trial, we extracted neural activity from 1 s before to 1 s after stimulus onset and assigned binary labels (1 = puff, 0 = no puff). Within each trial we slid a 20-ms window in 10-ms steps. In every time window, we evaluated the average activity of each neuron at each trial. We used the standard LIBSVM toolkit^45^ and fed the SVM module with neural activity and binary label. Training and testing were performed using a 10-fold cross-validation procedure. Data were divided into 10 equally sized and disjoint sets; in each fold, one of the sets (10% of the data) was used for testing, and the remaining 9 sets (90% of the data) were used for training the classifier. In each fold, the SVM regularization parameter was determined using a grid-search optimization using another (internal) 10-fold partition of the training data. Overall, the 10-fold cross-validation testing process resulted in 10 accuracy rates per time bin.

For k-means clustering in Figs. 3q and 3r, we divided the temporal traces into two stages (during air puff stimulus and without stimulus), and used the standard deviations of spike rate and subthreshold electrical activity during corresponding stages as features for each neuron. PCA method is first used to reduce the feature dimensionality from 4 to 3. We then used k-means method to separate neurons into four different clusters based on the features, after which the clustered groups of neurons can be precisely distinguished.

### *In vivo* vascular imaging and blood flow velocity measurement

Two wild-type mice were used for vascular imaging. Both were briefly anesthetized with isoflurane before retro-orbital injection with fluorescent solutions (200 μL each, 5% w/v): one mouse received 70-kDa dextran-conjugated Texas Red, and the other was injected with 70-kDa dextran-conjugated Rhodamine B. The mice were then head-fixed under the objective lens, and kilohertz intensity images were recorded for blood flow velocity measurement, while half-FOV images were recorded for calculating and comparing different fluorescence lifetimes.

Structural images were acquired for 5 s using the TPVI system at a frame rate of 916 Hz and a post-objective power of 200 mW via a 25× Olympus objective. First, to better analyze the temporal changes in blood flow and fluorescence lifetime, the raw vascular imaging data were denoised using the SUPPORT algorithm^46^ to eliminate random shot noise. The denoising model was trained using the raw data acquired by HS2PM, while its hyperparameters adopt the default parameters of the SUPPORT algorithm. Vessel segments were then manually traced over the average intensity of the recorded structural images (150 μm below pial surface), and the kymographs of the vessels were plotted for blood flow velocity calculation. Kymographs were generated by plotting pixels along the traced blood vessels (horizontal axis) over time (vertical axis), with dark diagonal streaks resulting from red blood cells (RBCs) moving along the vessel segment. An automated method developed by Chhatbar and Kara was used to measure the slope of RBC streaks using Sobel filtering and iterative Radon transforms^47^, and then the blood flow velocity of each segment was calculated every 92 frames (∼0.1 s). The diameter of the vessel segment was measured by extracting the vessel boundary: pixel brightness along a line orthogonal to the vessel’s length was extracted, and the diameter was determined as the distance between the two edges detected by a Radon transform.

## Resource Availability

### Lead contact

Further information and requests for resources and reagents should be directed to and will be fulfilled by the lead contact, Bo Li.

### Materials availability

This study did not generate new unique reagents.

### Data and code availability

All data generated in this study are included in the published article. Datasets and codes are available from the corresponding author upon reasonable request.

## Acknowledgements

This work was supported by the National Natural Science Foundation of China (T2222006, 32471142, 32400928), China Postdoctoral Science Foundation (BX20230081), Science and Technology Commission of shanghai Municipality (20JC1419500). The authors thank Michael Z. Lin (Stanford University) for the generous gift of the ASAP5 voltage indicator. The authors also thank Yichi Su for assistance with plasmid amplification, verification, and coordination of subsequent AAV packaging.

## Author Contributions

Conceptualization: B.L.; methodology: B.L., M.Z., S.P.L., C.H.K, H.Y.; biological experiments: Y.G.Z., Y.F.Z., X.Y.G., H.Y.; Zemax optical simulation: M.Z., J.H.H., F.X.; investigation: B.L., M.Z., S.P.L., H.Y.; visualization: M.Z., S.P.L.; funding acquisition: B.L., Y.M., L.C., M.Z.; project administration: B.L.; writing: B.L, M.Z., S.P.L., H.Y., Y.G.Z., Y.M., L.C., J.C.W.

---

**Extended Data Fig. 1.**
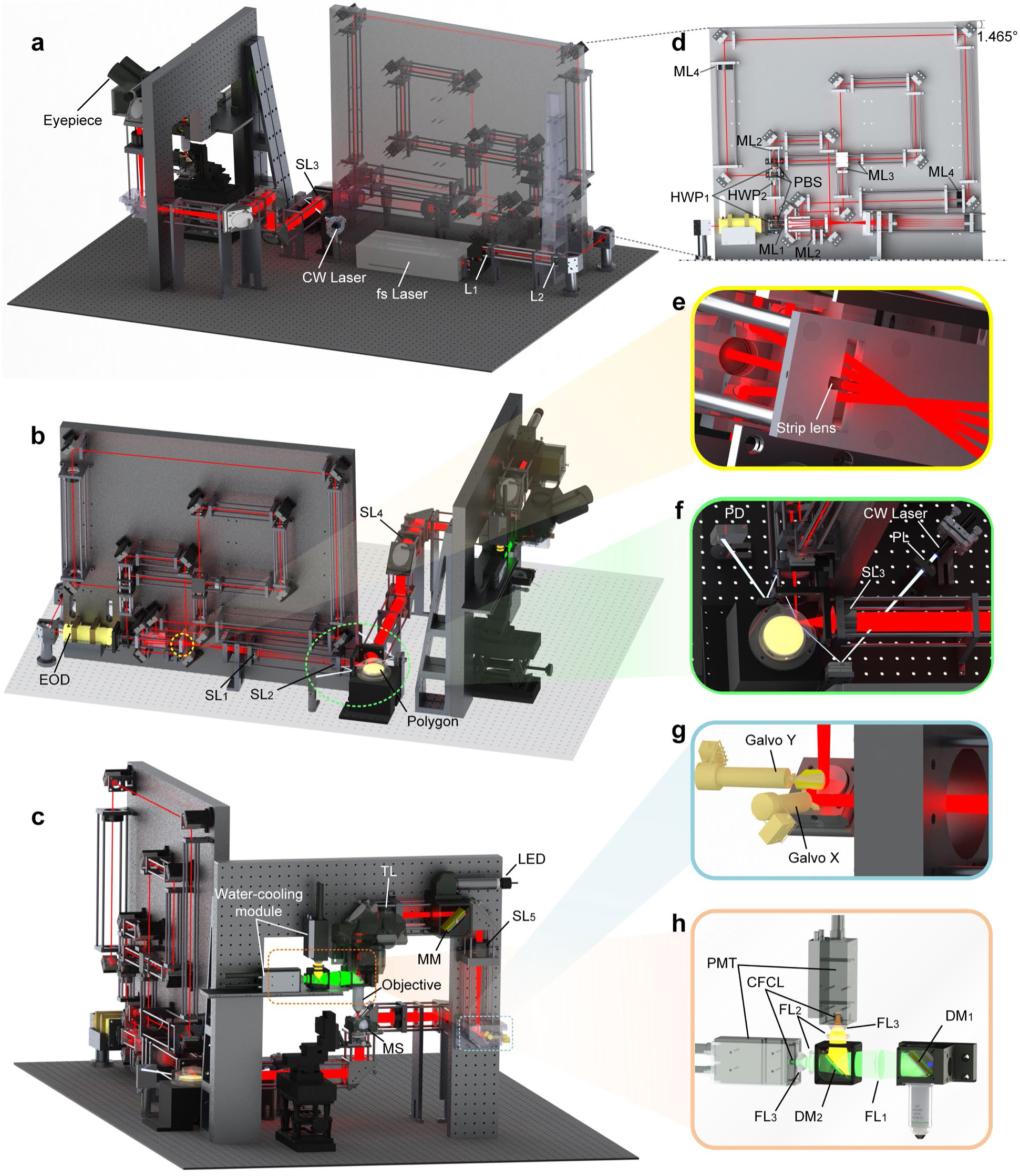
Opto-mechanical design of HS2PM. **a-c**, SolidWorks renderings of the complete HS2PM system, illustrating integration of optical and mechanical components. Excitation and fluorescence collection beam paths are shown. Insets highlight critical subsystems. **d**, Multiplexing module. The femtosecond laser is split into four beams using three HWPs and PBSs, with each beam routed through a distinct relay lens pair (ML_1_-ML_4_) and launched at a unique angle. **e**, Custom strip achromatic lens for channel 1, machined on both sides to prevent obstruction of adjacent beams. **f**, Start-of-line (SOL) signal detection: a 780 nm continuous-wave laser is focused onto the polygon scanner via a plano-convex lens (PL), and reflected signals are captured by a photodiode to generate TTL sync pulses per facet transition. **g**, Galvanometer scanner assembly for bidirectional *y*-axis scanning. **h**, Dual-channel fluorescence detection system. EOD, electro-optic deflector; SL, scan lens; TL, tube lens; DM, dichroic mirror; MM, movable mirror; MS, mouse stage; LED, light emitting diode; FL, fluorescence-collection lens; CFCL, custom frustum-of-cone light guide; PMT, photomultiplier tube.

**Extended Data Fig. 2.**
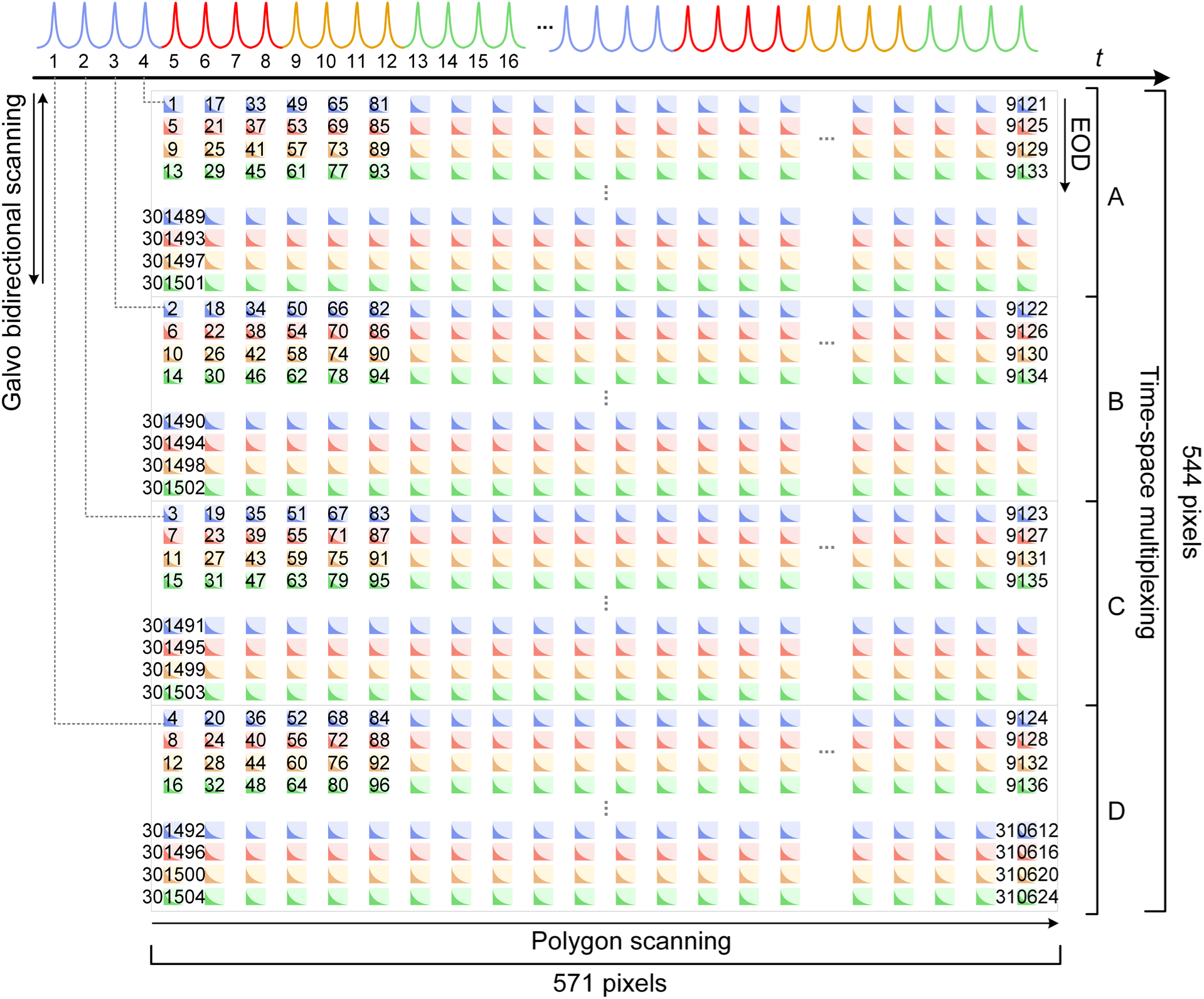
Schematic of pixel-wise order map of each frame.

**Extended Data Fig. 3.**
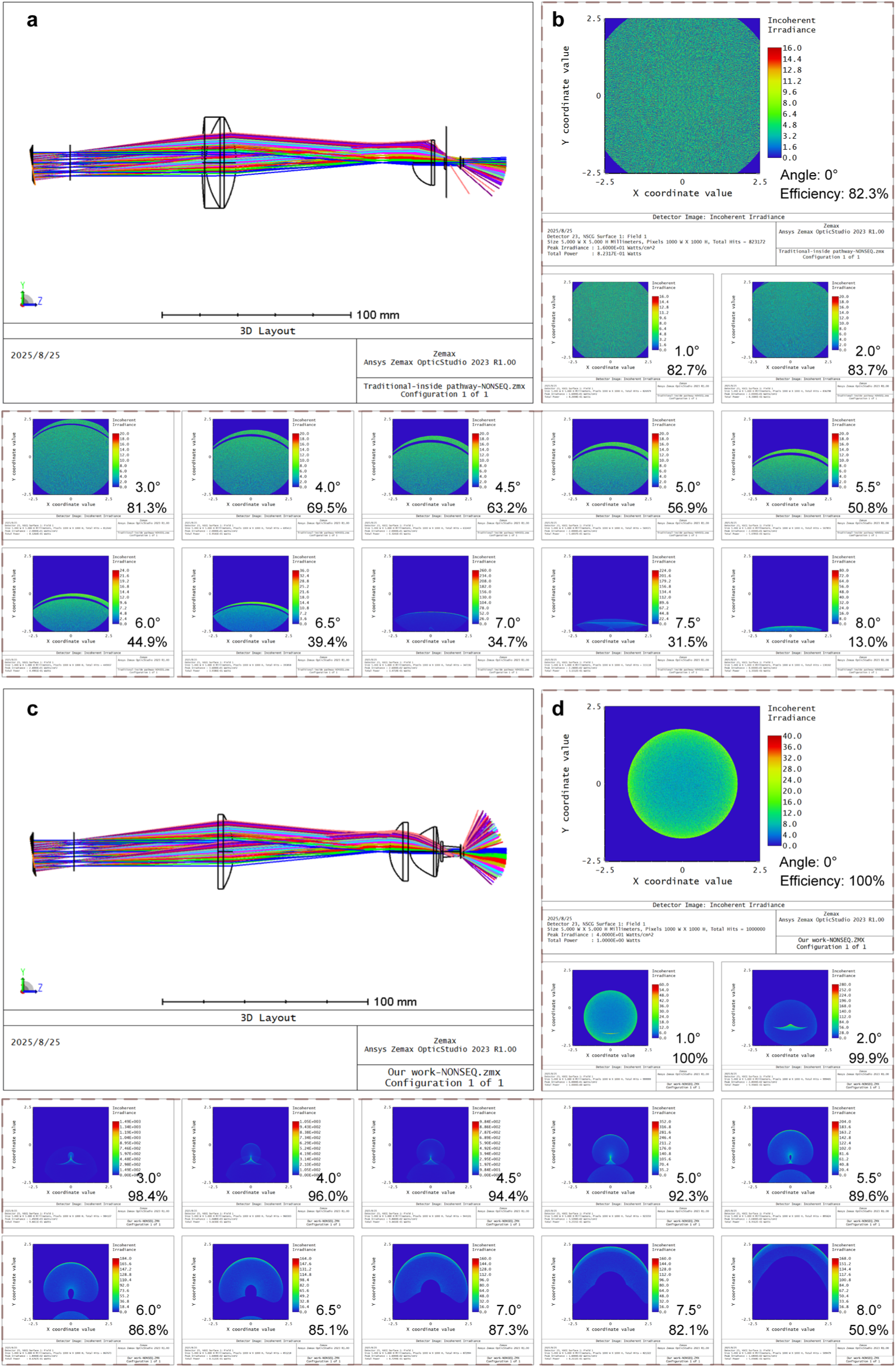
Comparison of fluorescence collection systems between conventional and the optimized design via Zemax optical simulations. **a**, Schematic 3D layout of a conventional fluorescence collection system simulated in Zemax. **b**, Simulated collection efficiency of the conventional fluorescence collection system across different angles. Each angle’s efficiency is defined as the ratio of power received by the detector to the initial 1 W input. **c**, 3D Zemax layout of the optimized fluorescence collection system. **d**, Simulated collection efficiency of the optimized system across different angles.

**Extended Data Fig. 4.**
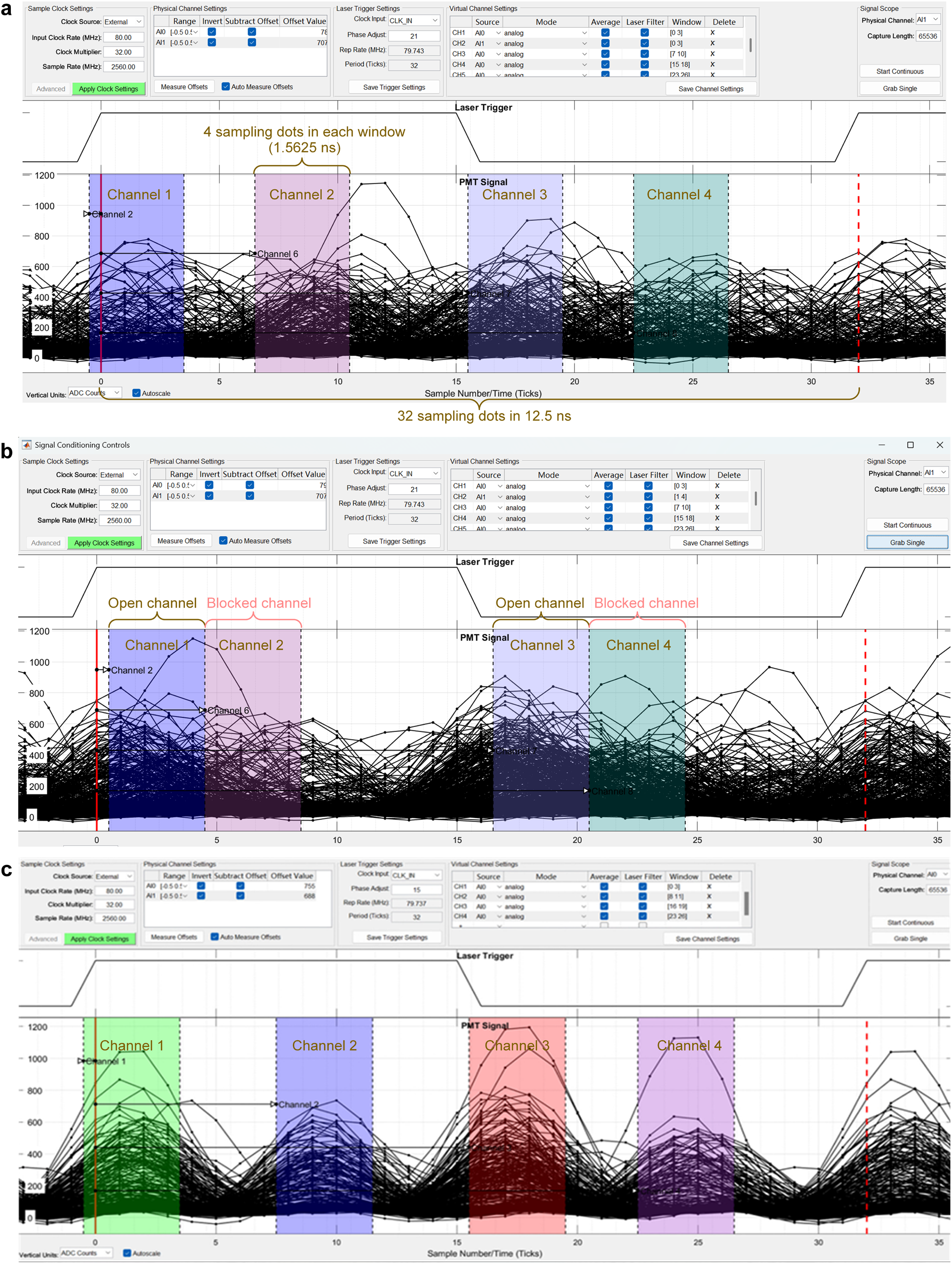
Data acquisition settings and the collected signals of different imaging modes. **a**, HS2PM, **b**, HS-FLIM, and **c**, HS2PM recorded free of cross-talk signals of *in vivo* autofluorescence.

**Extended Data Fig. 5.**
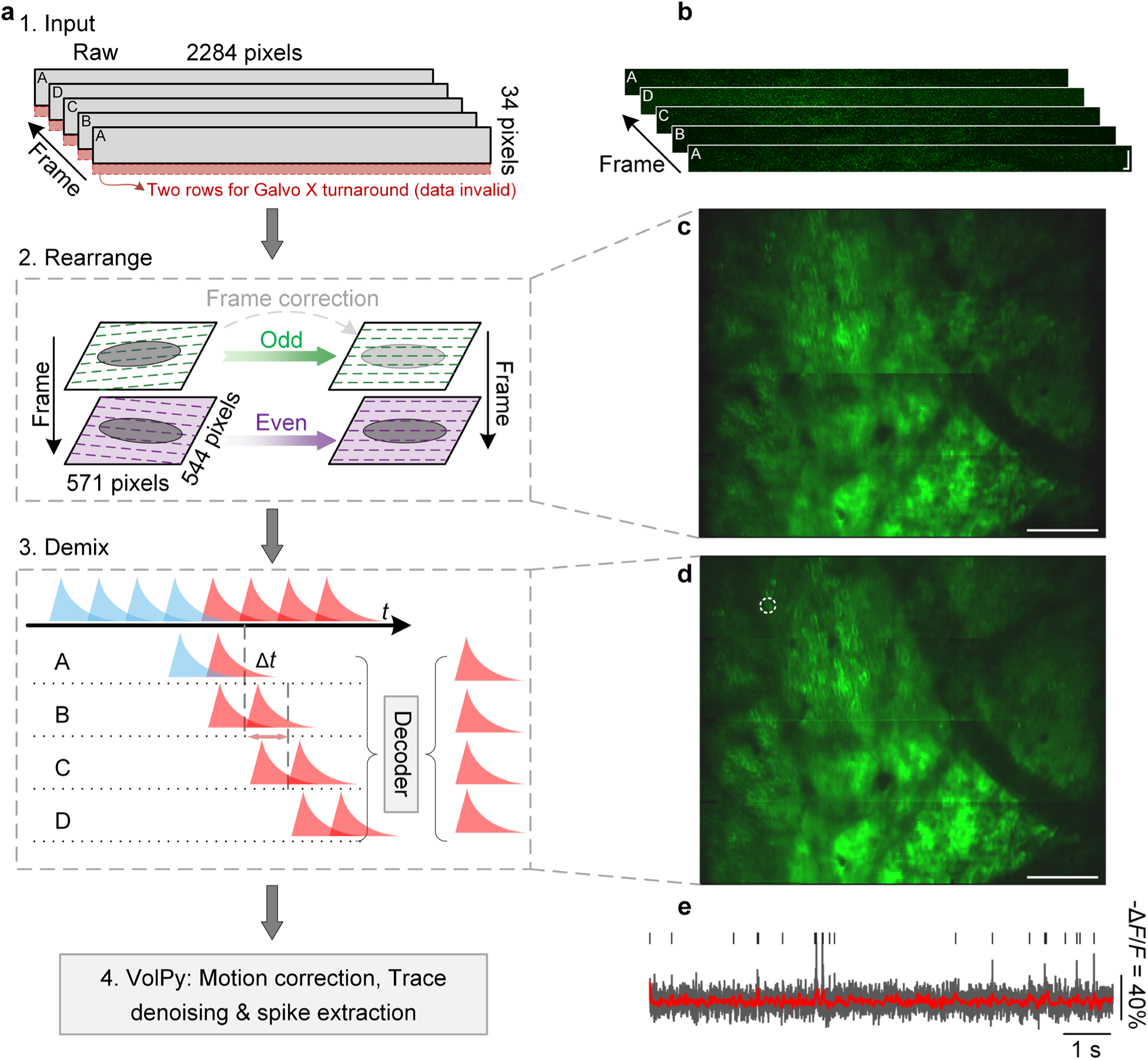
Computational pipeline for HS2PM voltage data extraction. **a**, Overview of the data-processing workflow for extracting neuronal voltage signals from raw imaging data. **b**, Representative raw frame (2,284 × 36 pixels). Only 34 rows are effective per frame, as 2 polygon facets are reserved for glavo turnaround during bidirectional scanning. **c**, Reconstructed raster-scanned image (571 × 544 pixels) after pixel rearrangement and odd-even frame correction. **d**, STDM demixing applied to **c** removes crosstalk across temporally multiplexed channels. **e**, Example -Δ*F*/*F* voltage trace (dark gray) extracted from a neuron identified in **d**. Red trace: subthreshold -Δ*F*/*F*; black ticks: detected spikes.

**Extended Data Fig. 6.**
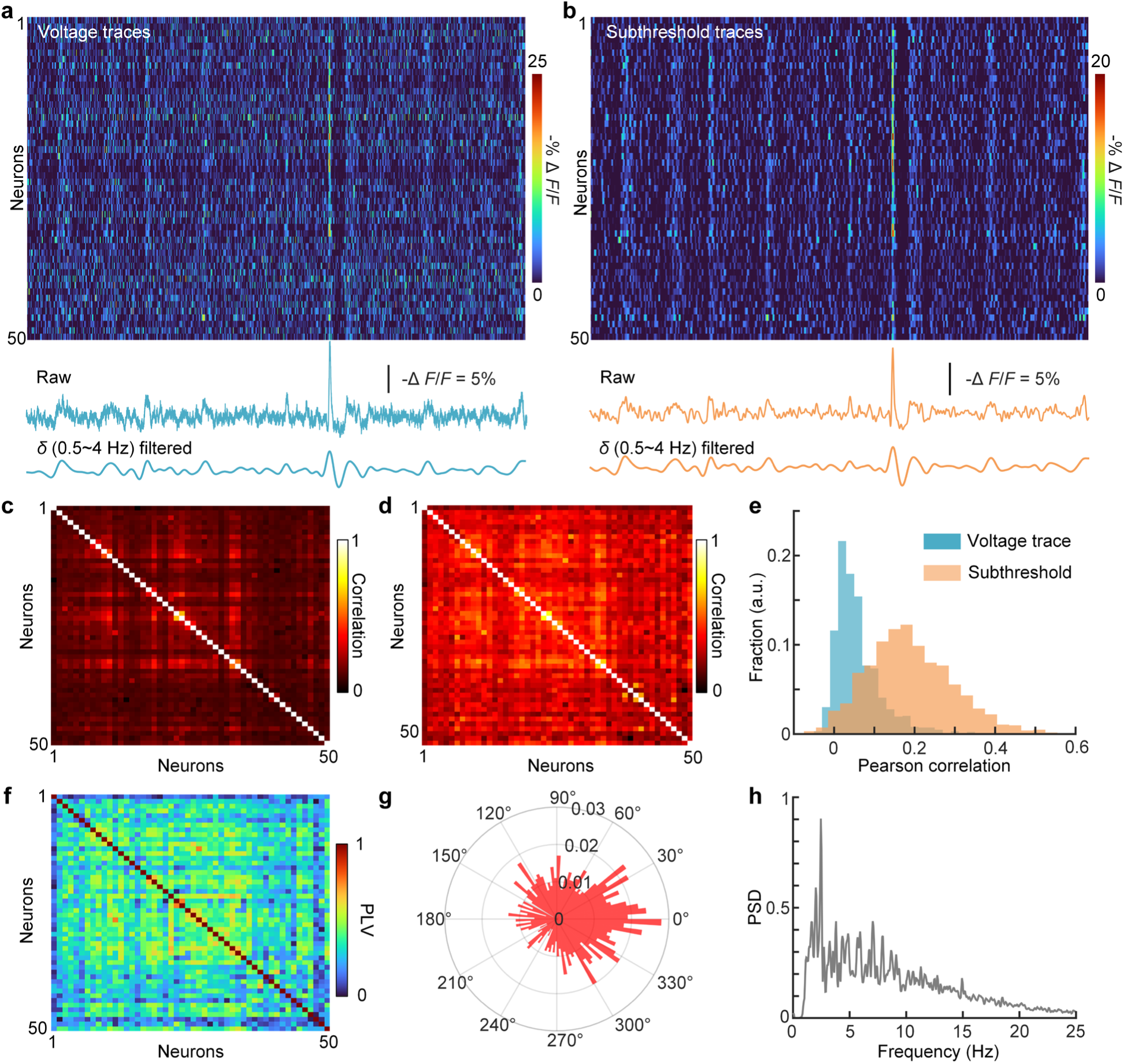
*In vivo* characterization of neuronal ensemble oscillations at 150 μm depth (related to Fig. 2f). **a-b**, Voltage traces and low-pass filtered subthreshold traces from 50 neurons over 10 s. From top to bottom: Heatmap of traces, averaged traces before and after filtered in the *δ* band (0.5-4 Hz). **c-d**, Cross-correlation matrices of voltage signals and subthreshold activities across all neurons. **e**, Distribution of Pearson’s correlation coefficients for spike and subthreshold signals. **f**, Phase-locking value (PLV) matrix among neurons. **g**, Polar plots of spike phase distribution relative to the 0.5-4 Hz membrane potential oscillations across neurons. **f**, Population power spectral density (PSD) of neuronal voltages activities in **a**.

**Extended Data Fig. 7.**
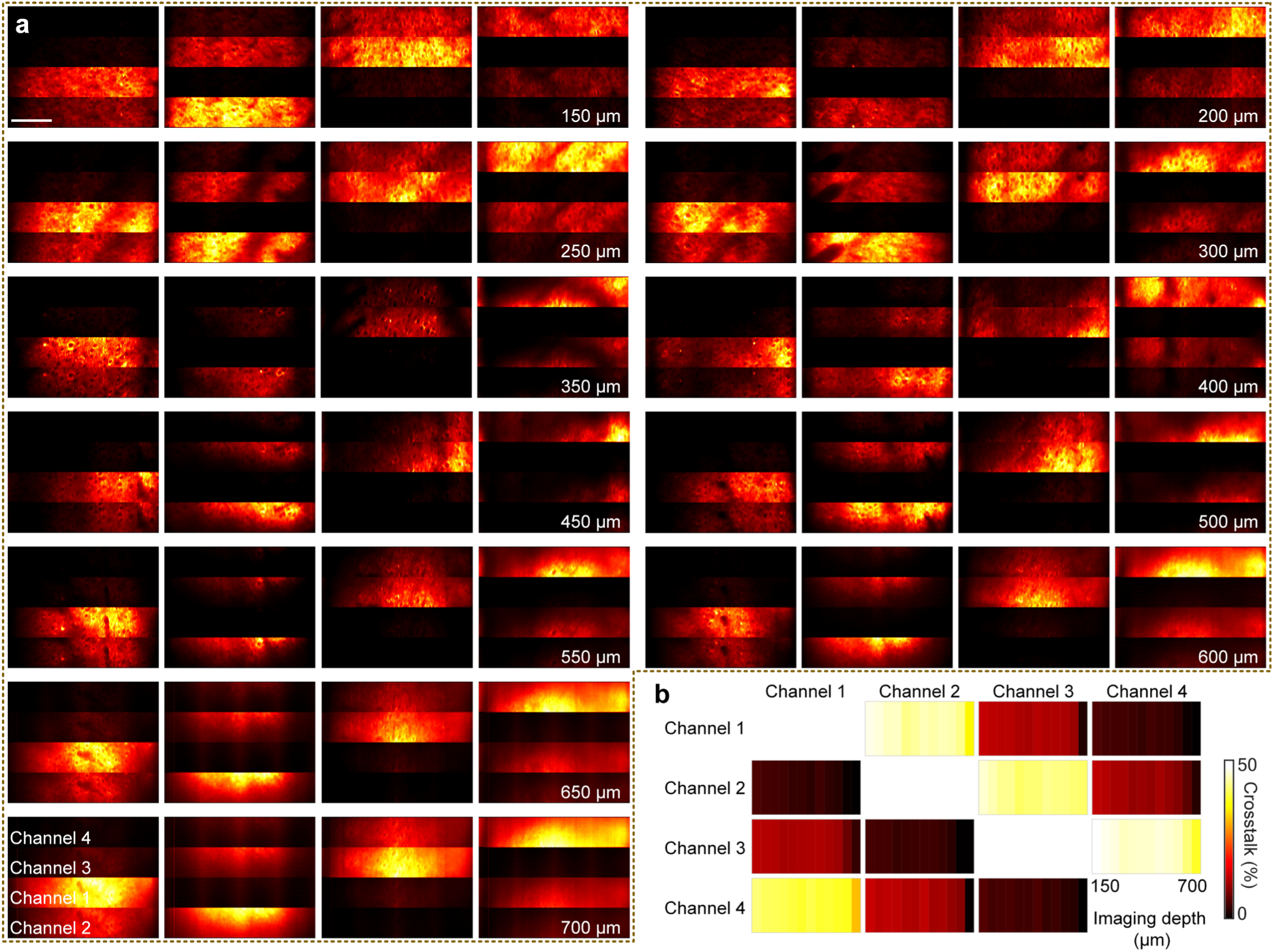
*In vivo* characterization of multiplexing channel crosstalk across cortical depths. **a**, Depth-resolved images (150-700 µm, in 50 µm steps) of four multiplexed channels. At each depth, four panels show signals detected by all four channels when only a single corresponding channel was unblocked for excitation. **b**, Crosstalk ratios among the four time-division channels as a function of depth, computed from the images shown in **a**. Scale bar: 100 μm.

**Extended Data Fig. 8.**
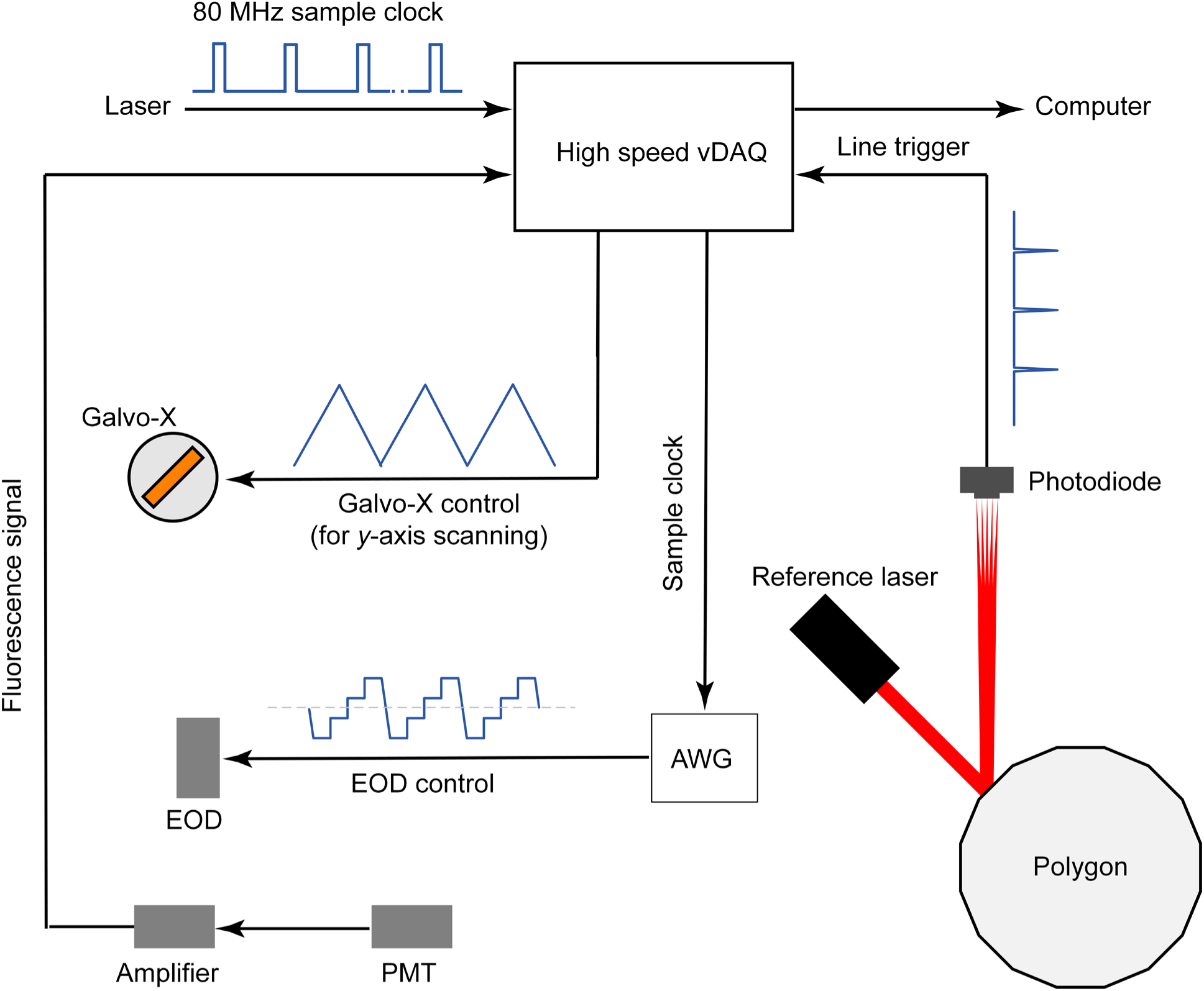
Synchronization diagram of the scanning and data acquisitions system. DAQ, data acquisition; AWG, arbitrary waveform generator.

